# Modelling pairwise and coalitional contests with reinforcement learning

**DOI:** 10.64898/2026.09.23.753778

**Authors:** Olof Leimar, Redouan Bshary

**Affiliations:** Department of Zoology, Stockholm University, 106 91 Stockholm, Sweden; Institute of Biology, University of Neuchâtel, Neuchâtel, Switzerland

**Keywords:** game theory, aggressive behaviour, territoriality, dear enemies, assessment

## Abstract

Game theory in biology started as an attempt to model animal contests by making assumptions about the fitness costs individuals pay in aggressive interactions. Among the different approaches, a notable one is to construct mechanistic models, using assumptions about behaviours and cognitive processes. Reinforcement learning is an important cognitive process, and we use it here to model pairwise and coalitional contests. We study situations where territorial neighbours become so-called dear enemies. A defending individual can get help from a neighbour through a defender-neighbour coalition against a challenger attempting territory takeover. In our model, coalition members have an advantage in contests in terms of costs of aggression, with the challenger being exposed to aggression from both members and each member only receiving part of the aggression from the challenger. We find that notable costs of border conflicts between neighbours, together with substantial advantages for coalitions, favour intervention. Neighbours intervene when they can help weaker defenders, in this way avoiding costs of border renegotiation with a new and potentially stronger territory owner. We also introduce different forms of perceived costs into the model, which we show correspond to the range from pure mutual assessment to partial self-assessment.

## Introduction

Game theory has long been used as a tool to study animal contests, starting from pioneering works in the 1970s and early 1980s [1, 2, 3]. Many game-theory models of contests are non-mechanistic, in the sense of making assumptions about the fitness consequences of different strategies meeting each other without specifying how the fitness consequences come about. There are also alternative approaches, and a promising one is to implement concrete behavioural mechanisms [4], in this way specifying the costs incurred during the progress of a contest over time. An essential component of such mechanisms could be reinforcement learning, which can operate as a form of assessment and has been used to model dominance hierarchy formation through pairwise contests [4, 5, 6, 7]. Our aim here is to extend the approach, by applying it to both pairwise and coalitional (two-against-one) contests, and by introducing more realism to the type of assessment processes [8] that direct learning and thus influence the outcome and duration of a contest.

Although most animal contests are pairwise, with one individual challenging another for a resource or a dominance position, there are also coalitional contests, the most common being a coalition of two against one. Coalitional contests occur both for territorial interactions, where a territory holder is challenged and gets help from a neighbour to defend its territory [9, 10, 11, 12, 13], and for interactions in dominance hierarchies, mainly in primate species [14, 15, 16, 17, 18, 19].

Here we focus on territorial interactions, where coalition formation depends on the so-called dear enemy effect [9]. Dear enemies are territorial neighbours who have settled any boundary disputes between them. If a territory is taken over from one of them by a new individual, there can be a cost for the neighbour, because of renewed and possibly costly border disputes with the new owner. In particular, if a relatively weaker territory holder is replaced by a stronger challenger, a neighbour could then face and perhaps lose a costly border contest. An alternative for the neighbour is to help in defending against a takeover by the challenger.

A main topic of interest for experimentalists investigating the phenomenon has been the circumstances that cause territorial neighbours to intervene and help in defending against a challenger. The questions include how the fighting abilities of challenger, defender, and neighbour influence the likelihood of intervention, and also whether intervention occurs at all in a given population or species. We use game-theory modelling to study these questions. We examine the influence of fighting abilities on intervention, and we also test three hypotheses about the effects of additional factors, inspired by previous work. The first hypothesis is that a neighbour might intervene primarily when the fitness costs of fighting are sufficiently low. As discussed in [13], if a challenger is strong and fighting is risky, the neighbour might avoid getting involved in a contest. A second hypothesis is that, if a neighbour gives help in order to avoid the cost of renegotiating its territory border with a winning challenger, the neighbour should intervene primarily when the costs of border negotiation are noticeable. The idea is inspired by field observations on rock pipits [10], in which it was found that interventions occurred on an island with tightly packed territories, but did not occur on the mainland, where the distance between territorial neighbours was greater. As the spacing or territories is likely to vary among populations and species, this could be an important factor explaining when dear enemy coalitions occur. A third hypothesis is that a sufficiently high fighting advantage for coalition members, for instance from coordinated attacks, promotes neighbour intervention.

Different types of assessment processes in contests have been analysed and discussed [8, 20, 21, 22, 23, 24], including mutual and self assessment. An advantage of implementing reinforcement learning in contest models is that such assessment processes can appear as variants of a general learning mechanism. Perceived rewards are basic elements of reinforcement learning, and for contest models these are expressed as internally generated rewards of performing aggressive behaviour (aggressive motivation) minus a perceived cost or penalty of being exposed to aggression from an opponent [4, 5]. The perceived costs depend on the contestants’ fighting abilities. If the costs depend only on the difference in fighting abilities, we refer to this as unbiased costs, and if there is a stronger dependence on the individual’s own fighting ability, the costs are self-biased. We use both self-biased and unbiased costs in our models, and these cost variants correspond to different assessment processes. Our aim is to achieve qualitative agreement with field observations for species with dear enemy coalitions, including some fiddler crabs [11] for which deviations from pure mutual assessment have been found [25].

After presenting the model and results, we discuss our current understanding of dear enemy coalitions, including questions about the relative involvement of each of the coalition members in a contest and about the adaptive interpretation of the phenomenon. Dear enemy coalitions are relatively rare among territorial species, which raises the question of whether neighbour intervention has evolved for the purpose of giving aid, or if it might be a form of spin-off from a general tendency to direct aggression against perceived threats. We also sketch our perspective on variation in assessment processes and suggest how further progress in understanding this phenomenon could be achieved.

### The model

The model describes a situation where an individual *i* challenges a defending territory holder *j* in a take-over attempt. There is a territory neighbour *k* who can choose to join the defender *j* in a coalition against the challenger *i*, in which case there is a coalitional contest. There can also be pairwise contests, either as border disputes between territory neighbours or as contests over territory ownership, for instance between a challenger *i* and a defender *j* if the neighbour *k* does not intervene. Here we explain the conceptual aspects of the model, with further details, including equations, given in the Supplements.

### Order of events

The sequence of events is, first, a border contest between the territory holders *j* and *k*, which is represented as a pairwise contest over a resource value *V*_b_ that is considerably smaller than the mean resource value *V*_t_ associated with possession of a territory. Next, individual *i* challenges the defender *j*, and the neighbour *k* then decides whether to intervene. With intervention, there is a coalitional contest of *i* against *j* and *k*, and otherwise there is a pairwise territorial contest between *i* and *j*. In either case, if the challenger wins and takes over the territory, there is a border contest between *i* and *k*, again as a pairwise contest over a resource value *V*_b_.

### Reinforcement learning

Our aim for the modelling is to use a simple implementation of reinforcement learning, to allow for evolved effects of the different positions in coalitional contests without introducing a great number of genetically determined traits. To achieve this we use action-value learning, inspired by the description in [26], but otherwise we follow previous reinforcement-learning approaches to contest modelling, which have used actor-critic learning [4, 5, 6, 7]. As illustrated in the Supplements, the evolutionary outcomes for models of pairwise contests with action-value learning and unbiased costs are qualitatively similar to ESSs for the sequential assessment game [27], and are also qualitatively similar to the corresponding evolutionary outcomes for models with actor-critic learning.

### Perceived costs, rewards, and action values

For coalitional contests, our approach is to represent one round of fighting as, for each of the coalition members, a round against a challenger where the challenger’s ability to inflict damage on a coalition member is reduced, and for the challenger as a round against the combined entity of the coalition members, having a greater ability to inflict damage on the challenger. Figure 1 depicts the perceived costs of a round of fighting, both for pairwise and for coalitional contests, and gives examples of the progression of action-value learning over time.

**Figure 1:**
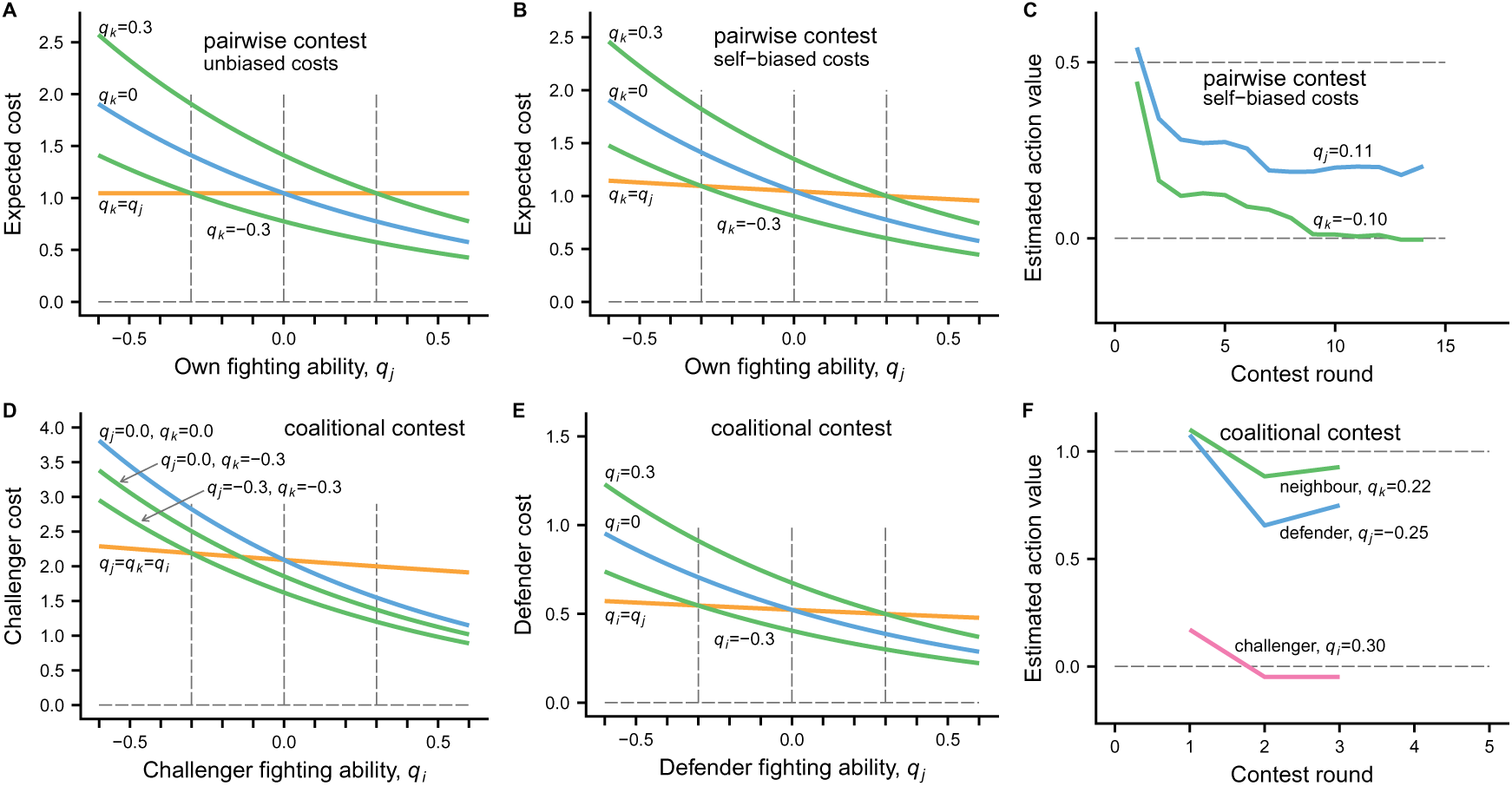
Illustration of basic model features. The expected perceived costs or penalties of a round of fighting are shown in panels (A) and (B), for pairwise contests between individuals *j* and *k*, as a function of the fighting ability *q_j_* for different values of *q_k_*. In (A), the perceived costs are unbiased, in being functions only of the difference *q_j_ − q_k_*, and in (B) they are self-biased, in the sense that there is a stronger dependence on an individual’s own *q_j_*, such that costs tend to be lower for higher fighting abilities, as illustrated by the curve for matched individuals (*q_k_* = *q_j_*; orange curve). The estimated value of performing an aggressive action is then an internal, perceived reward *v_j_*minus the perceived cost. Panel (C) shows these estimated action values for a pairwise contest with the self-biased costs from (B). Individual *k* gives up in round 14, having estimated a negative action value. Panels (D), (E), and (F) show the corresponding information for coalitional contests between a challenger *i* and a coalition of a defender *j* and a helping neighbour *k*. The costs now depend on all three fighting abilities *q_i_*, *q_j_*, and *q_k_*, and the single challenger *i* has a disadvantage in perceiving higher costs (D) than the coalition members, such as the defender *j* (E; note that the scales on the *y*-axes differ). In the coalitional contest in (F), the challenger *i* gives up already in round 3, in spite of having a higher fighting ability than both *j* and *k*.

An individual’s fighting ability (e.g., *q_j_*) expresses the ability to inflict costs on opponents and could, for instance, be the logarithm of body size. For the model we assume that fighting ability has a normal distribution in the population, with mean zero and standard deviation *σ_q_*, with *σ_q_* = 0.3 for the analyses in the main text. Mean zero fighting ability could be achieved, for instance, by choosing a suitable scale for body size.

The costs per pairwise fighting round in Fig. 1 are exponential functions of the difference *q_j_ − a*_1_*q_k_* (plus random noise), with *a*_1_ = 1.0 for the unbiased costs in panel A and *a*_1_ = 0.85 for the self-biased costs in panel B. A self-biased cost for individual *j* means that the cost decreases more rapidly with increasing *q_j_* than it increases with increasing *q_k_*. For matched opponents (*q_j_* = *q_k_*), this means that the perceived cost per fighting round is lower for higher fighting abilities, as shown by the orange curve in Fig. 1B. In addition to the costs, we assume that an individual perceives an internal reward (e.g., *v_i_*) from performing aggression, which we interpret as aggressive motivation or, using an animal psychology concept, as a primary reward. This reward is assumed to be genetically determined. The net perceived action value is then the perceived reward minus the perceived cost. An example of a pairwise action-value learning contest is shown in Fig. 1C, where individuals use their perceived net rewards to update an action-value estimate, and choose the aggressive action as long as the estimate is positive. The starting value of the estimate is genetically determined. In the example in Fig. 1C, the starting values are fairly high, which can be seen as a form of exploration, in the sense that a weaker contestant typically learns about the action-value over at least a few rounds before giving up.

For the challenger *i* in a coalitional contest, the cost per fighting round in Fig. 1D is simply the sum of the costs that would apply in pairwise contests with each of the coalition members. For the coalition members, such as the defender *j* in Fig. 1E, the cost is instead half what it would be when meeting *i* in a pairwise contest. The idea behind this is that, supposing the contestants strike blows against each other, the challenger would receive the sum of blows from the two coalition members, whereas each of these members would receive only half of the blows from the challenger. This corresponds to having parameter values of *b*_1_ = 1 and *b*_2_ = 0.5 in the model, and gives the coalition a considerable advantage, which is illustrated by the coalitional contest in Fig. 1F. By changing these parameters, we can change the strength of the advantage for a coalition. For instance, having *b*_1_ = *b*_2_ could be seen as an approximation to a case where members alternate between rounds in confronting a challenger.

In our model, the starting values for estimates by the challenger, defender, and neighbour in a coalitional contest are influenced by genetically determined traits, one for each of these roles. For instance, in Fig. 1F the challenger starts at a lower estimated action value, causing this individual to more readily give up, whereas the defender has a higher starting value, and the helping neighbour is intermediate between these.

#### Intervention

A neighbour bases the decision of whether to intervene on the threshold condition that a helping liability is positive, and pays a small opportunity cost for temporarily leaving the home territory. The helping liability is a linear function of the neighbour’s action-value estimates for the challenger and the defender, and the parameters (intercept and two slopes) of the linear function are genetically determined traits. When deciding on intervention, the neighbour has experience from a border contest with the defender but there is no prior contest with the challenger. We assume that the neighbour still has some information about the challenger’s fighting ability from visual cues, for instance cues of size. For simplicity, we make the modelling assumption that a pair of individuals gain information about each other corresponding to ‘a fraction of a fighting round’, thus influencing the action-value estimate. In this way the model gains an element of initial visual assessment, without needing to make special assumptions about such a process.

As an alternative to having a single condition for intervention, in the Supplements we also briefly discuss the case of having separate threshold conditions for the action-value estimates for the challenger *i* and the defender *j*. This is inspired by experimental results, such as [12], suggesting that there could be two conditions.

#### Evolutionary analysis

We use individual-based evolutionary simulations to analyse the model. For simplicity, individuals are assumed to be hermaphrodites, with the genetics of each trait being an independent implementation of the infinitesimal model of quantitative genetics [28]. There are 6000 individuals in a simulated population, split into groups of three (2000 groups), with random assignment to the roles of challenger, defender, and neighbour, and randomly determined fighting abilities. After one season of contest interaction, the next generation is formed by mating over the entire population (for simplicity), with the contributions from an individual being proportional to that individual’s accumulated fitness gains and losses. Fitness losses are determined by the accumulated fighting damage, with damage being proportional to the perceived costs. We have run simulations over very many generations, up to 2*×*10^6^, in order to get reasonable estimates of multi-trait evolutionary equilibria. There are traits for starting estimates, internal perceived rewards, a learning rate, and parameters for the neighbour’s helping liability. The model and simulation details are presented in the Supplements, including brief definitions and notation in Table S1 and examples of evolutionary trajectories in Fig. S1.

## Results

Figure 2 illustrates probabilities of winning and contest durations as functions of the fighting abilities for pairwise and coalitional contests. Panels A, B, and C in the figure correspond to the initial border contest between a defender *j* and a neighbour *k* for self-biased costs, and panels D and E illustrate a following coalitional contest, where the neighbour intervened. There is a pronounced advantage for the coalition, such that the challenger needs to have a considerably higher fighting ability than the coalition members in order to win (D), and the longest contests occur for these stronger challengers (E). Panel C illustrates effects of self-biased costs on the duration of pairwise contests between approximately matched opponents, with longer contests for stronger pairs. This can be compared with panel F, which shows the corresponding result for unbiased costs (the orange lines show linear regressions; highly significant in C and non-significant in F).

**Figure 2:**
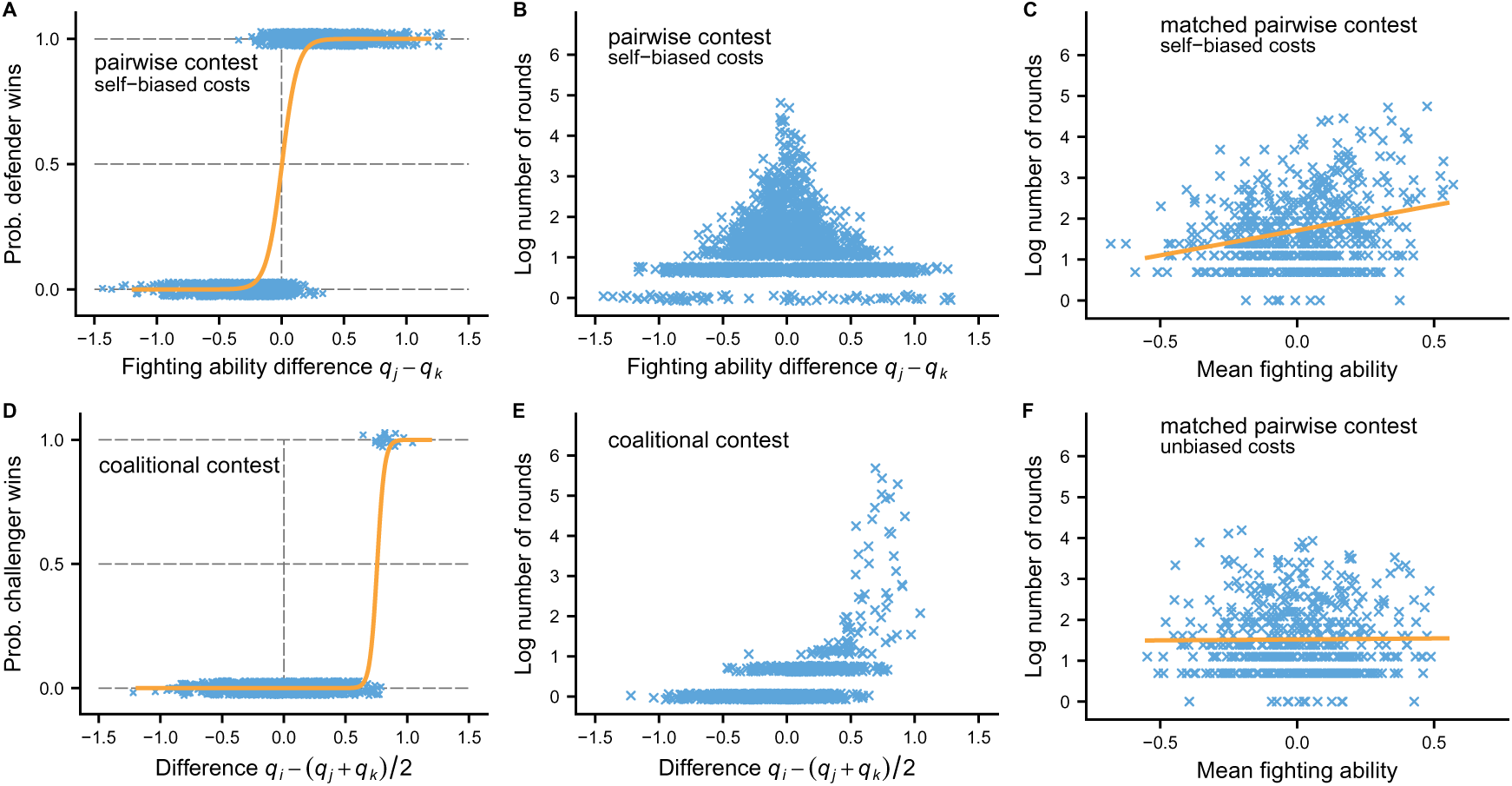
Basic characteristics of pairwise and coalitional contests at an evolutionary equilibrium. Comparing panels (A) and (D) shows that to win a pairwise contest a challenger *i* needs a smaller fighting ability advantage over the opponent than the advantage over the mean of opponents needed to win a coalitional contest. Similarly, comparing (B) and (E) shows that pairwise contests tend to be longer than contest contests. The longest coalitional contests occur for cases where the challenger *i* has around 50% chance of winning (D and E). Panels (C) and (F) show the distributions of the length of pairwise contests between approximately matched individuals (*|q_i_ − q_j_| <* 0.15), for self-biased costs in (C) and unbiased costs in (F). In (C), pairwise contests tend to increase in duration with the fighting abilities of matched opponents, as a consequence of the smaller costs perceived by stronger individuals.

Figures 3, 4, and 5 show results on the circumstances under which neighbour intervention is evolutionarily favoured. There is both the question of whether intervention occurs at all and, in situations where interventions sometimes occur, for which configurations of fighting abilities *q_i_*, *q_j_*, and *q_k_* intervention is more likely.

**Figure 3:**
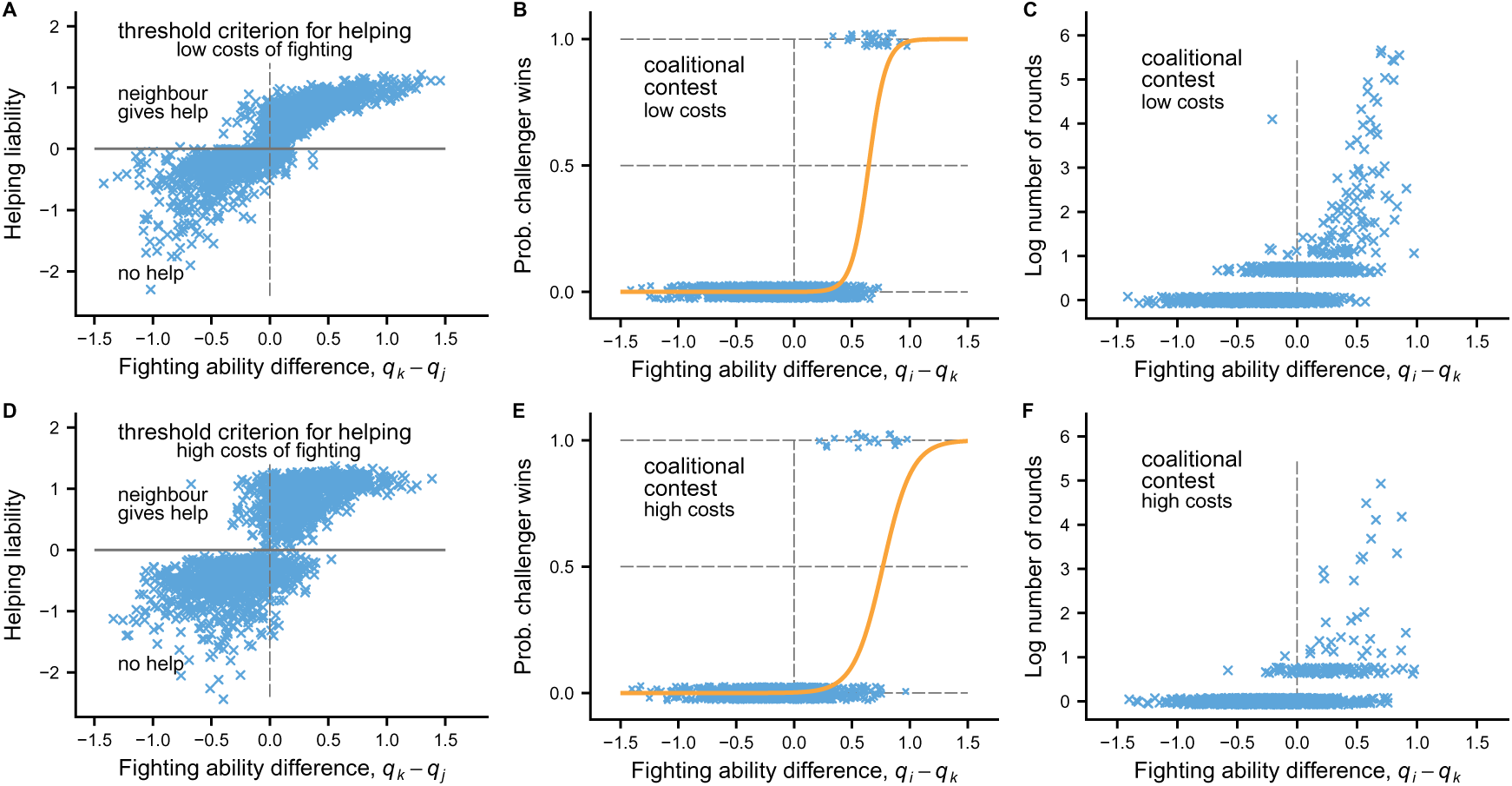
Illustration of the influence of the magnitude of fitness costs of fighting on the tendency of a neighbour *k* to intervene to help a defender *j* against a challenger *i*. The neighbour *k* uses a threshold criterion of a helping liability being positive for the decision of whether to help. The helping liability is a function of the action-value estimates obtained by *k* and is correlated with the difference in fighting ability between the neighbour *k* and the defender *j*. The top panels correspond to the situation in Fig. 2. Panel (A) illustrates that *k* often intervenes, and more often when the defender *j* is relatively weaker. Comparing with (D), where the fitness costs of fighting are four times higher, shows that the tendency to intervene seems to depend only weakly on the costs: there is 52% intervention in (A), and 46% in (D). Panels (B), (C) and (E), (F) show probabilities of the challenger *i* winning a coalitional contest (given that *k* intervenes) and the duration of such coalitional contests. Note that the probabilities of winning are similar for low and high costs, but that the durations of coalitional contests are substantially shorter for high costs.

We find that the magnitude of the fitness costs of fighting only have a weak effect on whether intervention occurs (Fig. 3A,D), although these costs influence the duration of contests, with shorter coalitional contests for higher fitness costs (Fig. 3C,F).

The magnitude of the fitness effects of boundary contests, on the other hand, can have a marked influence on intervention, and if these effects are small enough the neighbour never intervenes (Fig. 4A,D). Such situations are more favourable for a challenger, who will have a higher probability of winning a pairwise ownership contest with the defender, compared to winning a coalitional contest (Fig. 4B,E). Similarly, if the the coalitional cost advantage is small enough, the parameters for the helping liability evolves such that the neighbour never intervenes (Fig. 5A,D).

**Figure 4:**
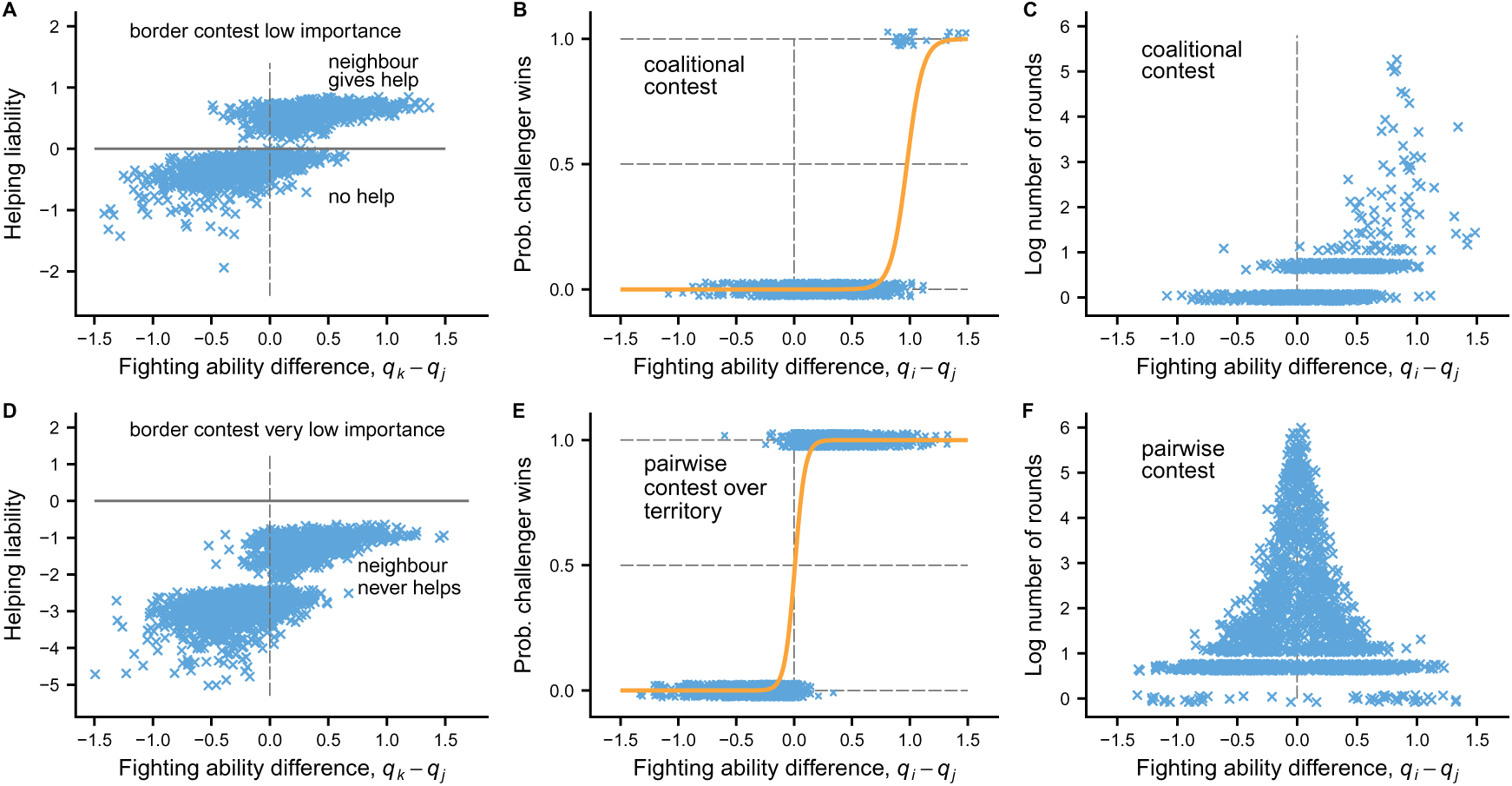
Similar to Fig. 3, but showing the influence of the fitness effects of boundary contests on coalition formation. In the top three panels, the fitness effects of boundary contests are smaller than in Fig. 3 (*V*_b_ = 0.10 vs. *V*_b_ = 0.25 in Fig. 3), and the neighbour *k* intervenes somewhat less often (43% intervention). The neighbour’s threshold criterion is based on a helping liability variable that is a function of the action-value estimates obtained by *k* and is correlated with the difference in fighting ability between the neighbour *k* and the defender *j*, just as in Fig. 3. In terms of winning and duration, the coalitional contests are similar to those in Fig. 3. For very small fitness effects of boundary contests (*V*_b_ = 0.02), as shown in panels (D), (E), and (F), the neighbour *k* never intervenes. Panels (E) and (F) show probabilities of the challenger *i* winning and the durations of the pairwise contests over ownership of the defender’s territory. Note that the challenger needs less of a fighting ability advantage to win such a pairwise contest, in comparison with winning a coalitional contest, and that the pairwise contests tend to be longer, with the longest contests occurring for approximately matched *i* and *j*.

**Figure 5:**
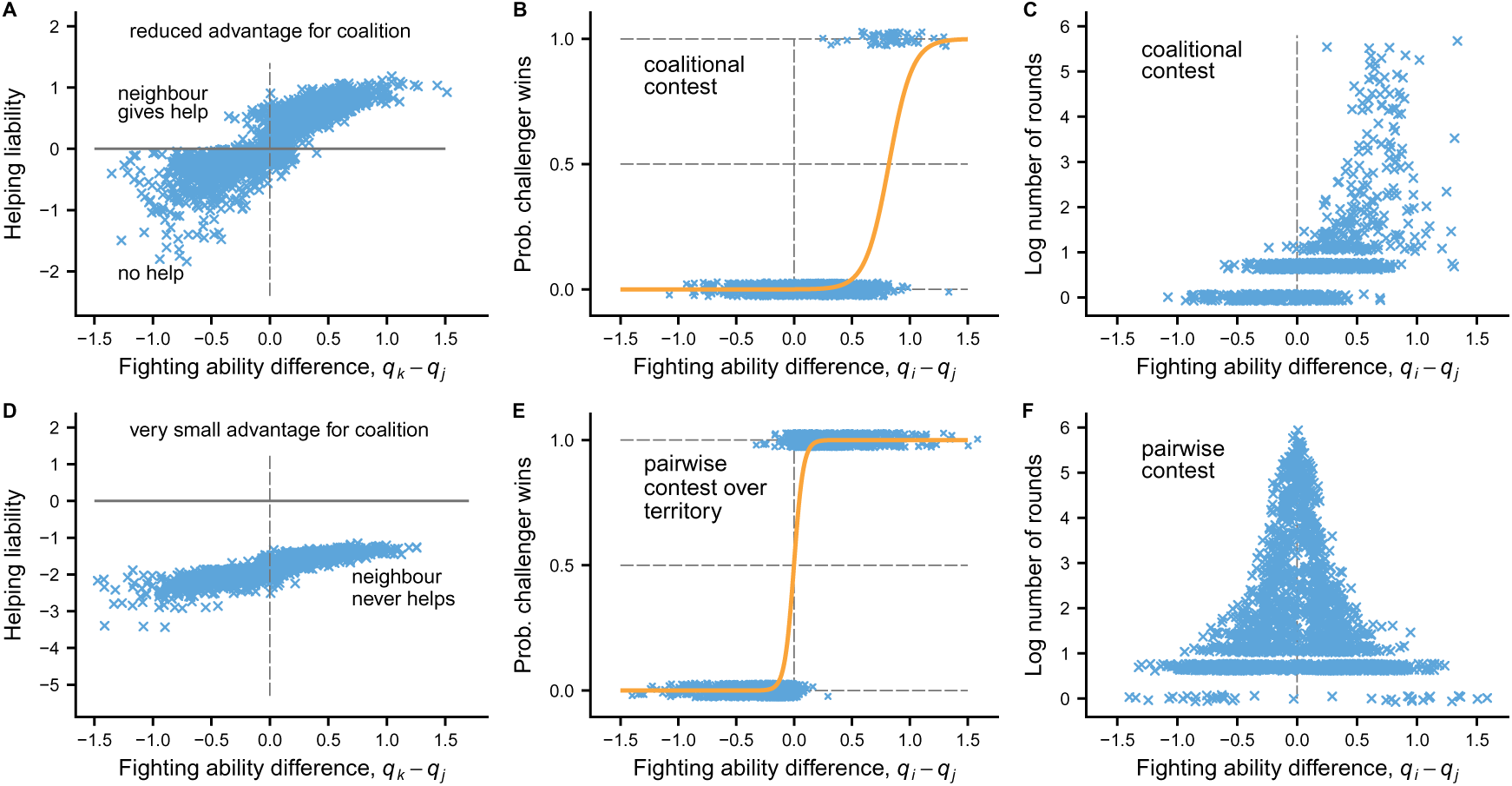
Similar to Fig. 3, but showing the influence of the cost advantage of coalition members *j* and *k* over the challenger *i* on neighbour intervention. For the top panels, with somewhat lower coalitional cost advantage than in Fig. 3 (*b*_1_ = 0.9*, b*_2_ = 0.6 vs. *b*_1_ = 1.0*, b*_2_ = 0.5 in Fig. 3), the neighbour’s threshold criterion for intervention is similar to that in Fig. 3A, and the rate of intervention (50%) is also similar, as are the other top panels, (B and C). Having *b*_1_ = *b*_2_ = 0.75 gives qualitatively similar results as in the top panels (not shown). In the three bottom panels the cost advantage of coalition members is very small (*b*_1_ = 0.6*, b*_2_ = 0.9, compared to a no-advantage case of *b*_1_ = 0.5*, b*_2_ = 1.0), and the neighbour *k* no longer intervenes to help the defender *j*. The pairwise contests between the challenger *i* and the defender *j* are similar to the pairwise contests in the corresponding panels of Fig. 4.

Concerning the configurations of fighting abilities, Fig. 3A,D, Fig. 4A, and Fig. 5A all show a pattern of two seeming clusters of cases where intervention either happens or not, with most interventions occurring when the neighbour has higher fighting ability than the defender (*q_k_ > q_j_*). The clusters correspond to a win or a loss by the neighbour *k* in the previous border contest with *j*. For instance, in 99% of the cases with intervention in Fig. 3A, the neighbour *k* had won the previous border contest with *j*. Thus, with our simple model implementation of border contests, with a discrete win/loss outcome instead of a graded adjustment of the border, the win vs. loss outcome of the border contest strongly influences whether intervention occurs.

Concerning the influence of the neighbour-challenger relative fighting ability on the decision to intervene, we find a weaker influence than that of the neighbourdefender relative fighting ability. This might to a large extent be explained by the limited information a neighbour has about the challenger at the time of making the intervention decision. Looking at the proportion of cases of neighbour intervention in which the challenger is relatively weaker (*q_i_ < q_k_*) we find it to be around 66% in each of Figs. 3A, 4A, and 5A, with the challenger being relatively stronger in the remaining cases.

Experiments sometimes investigate assessment by examining the relation between pairwise contest duration and the fighting ability of closely matched contestants [22], in a similar way as our model results shown in Fig. 2C,F. It is nevertheless more common to examine the relation between contest duration and fighting ability separately for winners and losers [22], and this is approach is inspired by [8]. Using our model simulations, we present results of this kind in the Supplements, for strongly and moderately self-biased costs as well as for unbiased costs (Figs. S5, S6). Our results agree qualitatively with the interpretations in [8], again illustrating that reinforcement learning can represent different kinds of assessment processes.

Examining contests between approximately matched opponents is of interest not only because it says something about assessment processes, but also because it can illustrate variation in the efficiency of different such processes in resolving a contest. For instance, for strongly self-biased costs, contests between approximately matched individuals of high fighting ability can become very long and result in pronounced fighting damage (Fig. S7). There would then be selection for reducing these costs, which could happen through the evolution of mechanisms that allow better assessment of relative fighting ability, possibly limiting the degree of self-biased assessment in contests.

## Discussion

We have shown that action-value learning [26] – arguably the simplest form of reinforcement learning – can be used as a cognitive mechanism in models of pairwise and coalitional contests (Figs. 1, 2). It is not to be expected that a single learning process could account for all aspects of animal contests, which show various behaviour patterns including threat signalling, but we find that models with action-value learning deliver results in reasonable agreement with other mechanistic approaches, such as the sequential assessment game [27] or models with actor-critic learning [4, 5]. Action-value learning can thus serve as a helpful ingredient in contest models.

When using action-value learning to investigate dear enemy coalitions in territorial systems, we find that neighbour intervention and coalitional contests evolve when the fitness effects of territory boundary conflicts are sufficiently important (Fig. 4) and when there is a notable contest advantage for a coalition of two against one (Fig. 5). Another main conclusion is that, if a territory holder is challenged by a floater individual, a territory neighbour intervenes and helps in defending against the challenger primarily when the neighbour is stronger than the territory holder (Figs. 3, 4, 5). The incentive for intervention can be understood by noting that it is a low-cost opportunity to prevent a challenger from taking over the territory and imposing larger costs through territory border conflicts with the neighbour. This conclusion is in accordance with previous suggestions [11, 12, 29, 30, 31], and means that the helping is asymmetric and driven by by-product benefits for the neighbour. Neighbours might intervene either to gain direct benefits or in order to induce future reciprocation, and these alternatives were put forward quite some time ago [9]. Subsequent observations indicate that the asymmetry makes reciprocation unlikely to occur for dear enemy coalitions [11, 12].

The weak influence of the challenger-neighbour (and the challenger-defender) relative fighting ability on intervention in our analysis is likely a consequence of our model assumption that a neighbour has limited information about a challenger. If there would be more information, and given the logic of our model, one might expect intervention to be more likely against a stronger challenger, because such an opponent would be particularly costly in a border conflict. Still, based on our simulations the evolved intervention decision appears to depend on the details of the costs for the different roles of challenger, defender, and neighbour.

There are both field observations [10, 11, 12] and experiments [12, 13] on intervention by dear enemy neighbours during territory challenges. For rock pipits [10], aggression by coalition members is described as sometimes alternating and some-times simultaneous, which might correspond to the situation in our Fig. 5A,B,C. For fiddler crabs, it is not clear precisely how aggressive interactions with intervention are structured. Some descriptions indicate that that both coalition members take part in a contest [11, 12], but there is also the statement that an intervening neighbour completely takes over the contest [30]. Our model can readily be modified to have different degrees of involvement of the roles of neighbour and defender in a coalitional contest, and we briefly discuss and illustrate this in the Supplements (see Fig. S8). For an extreme case where an intervening neighbour completely takes over the contest with a challenger, we find that there is selection against intervention. This makes sense as it should be preferable for the neighbour to encounter the challenger in a pairwise border dispute, which would be over lower stakes than a pairwise contest over the defenders territory. Overall, it appears that the question of whether dear enemy coalitions are adaptations to provide assistance in defending against a challenger or, at least to some extent, they should be interpreted as a side effect of a not so fine tuned adaptation for territory holders to treat any nearby challengers as a threat. Finally, we note that our current model is simplified in not taking into account possibilities such as choices by a challenger about whom to challenge [29].

A previous game-theory model of dear-enemy coalitions [31] differs from ours in several ways, for instance by not using specific assumptions about cognitive mechanisms such as reinforcement learning, but also shows some similarities. Among the similarities are basic assumptions of equal involvement by coalition members in a contest and the general conclusion that stronger neighbours tend to intervene in favour of weaker defenders.

We have focussed on dear-enemy coalitions in territorial systems, for which there is empirical work, but there is also a wealth of studies on coalitions in primate dominance hierarchies [e.g., 14, 15, 19, 16, 17, 18]. A qualitative difference is that such coalitions often or even typically involve the build-up of social bonds between members, which is why we have not dealt with them here. In order to analyse the phenomenon using mechanistic game-theory models, one would need to include reinforcement learning, as we have done here, together with an implementation of social-bond dynamics, for instance as in a previous model of helping in vampire bats [32].

The general idea that game theory in biology can profit from integrating function and mechanism into models has a history of at least a few decades [4, 24, 33, 34, 35, 36, 37], with both successes and challenges. Among the difficulties is that models with explicit mechanisms can be complex to construct and analyse, with multitrait individual-based evolutionary simulation of explicit life histories as a main method. There is on the other hand the crucial advantage of an opportunity for closer integration of theoretical modelling and observation, which can be beneficial to both. As examples from our work here, for coalitional two-against-one contests it could be helpful to have more information about how the coalition members behave in coordinating their aggressive behaviour and how the target of their aggression distributes aggressive behaviour between them.

For assessment processes, which have been much studied and discussed [8, 20, 23, 24, 25, 38, 39], variation in cognitive and other behavioural mechanisms could be a major explanation for the observed range from strictly mutual assessment to different degrees of self-biased assessment. There has been a tendency among empiricists to treat this variation as discrete and to postulate distinct so-called assessment strategies [8, 23]. This conceptualization has been criticized, with the proposed alternative that the observed variation corresponds to a continuous spectrum of mechanisms [21]. Our current results show that this alternative has promise (Fig. 3, Figs. S5, S6, S7). Making assumptions about the processes and mechanisms through which contestants regulate their behaviour allows for helpful comparisons between modelling results and observations. Further work is needed to make progress on the matter, but it seems clear that models that combine mechanistic elements with evolutionary analysis have the potential to contribute to the study of social behaviour.

## Code availability

C++ source code for the individual-based simulations is available at GitHub, together with instructions for compilation on a Linux operating system: https://github.com/oleimar/avcont1.

## Author contributions

OL and RB developed the conceptual aspects of the model; OL implemented and performed model analysis; OL wrote the manuscript; RB helped revise the manuscript.

## Supplementary information

### Supplementary tables

#### Model details

For completeness, we give a full description of the model here, repeating some parts of the description in the main text. The aim is to investigate a situation where a challenger *i* is attempting to take over the territory held by a defender *j*, and there is also a neighbour *k* who can decide to intervene and form a coalition with *j* against *i* (see Table S1 for notation and brief definitions). The indices *i, j, k* are used to conventionally denote these roles; we might have *i* = 1, *j* = 2, and *k* = 3. For the takeover attempt, the resource values (i.e., fitness differences between winning and losing) are *V_i_*, *V_j_*, and *V_k_* for the roles, with *V_i_* = *V_j_* = *V*_t_ *>* 0 and *V_k_* = 0. If the neighbour *k* does not intervene, the takeover attempt becomes a pairwise contest between *i* and *j*, and if *k* intervenes it becomes a coalitional contest between *i* and the coalition of *j* and *k*. If either *j* or *k* drops out of the coalitional contest, there is a continuation of the takeover attempt as a pairwise contest.

The model also takes into account border disputes between owners of nearby territories. The contested resource value for border disputes is assumed to be *V*_b_, with *V*_b_ considerably smaller than the mean value of a territory, *V*_t_. At some time before the takeover attempt, we assume that there has been a border contest between *j* and the neighbour *k*, after which they can be regarded as ‘dear enemies’. From this border contest, *j* and *k* gain information abut each other’s fighting abilities. Further, if the challenger *i* succeeds in taking over the territory from the defender *j*, there is a subsequent border contest between *i* and the neighbour *k*.

The neighbour *k* needs to decide whether to intervene in favour of the defender *j* without having any prior contest with *i*. We assume that *k* still has some information about the fighting ability of *i* from visual cues, for instance cues of size, before any contest between them. For simplicity, we make the modelling assumption that a pair of individuals gain information about each other corresponding to ‘a fraction of a fighting round’, meaning an update as in eqs. (S3, S4) below, but with a learning rate that is a fraction of the rate from a fighting round (in our simulations, we used a fraction of 0.5), and without any accumulation of fighting damage. We assume that a pair of individuals acquire this information when they first encounter each other. In this way the model gains an element of initial visual assessment, without needing to make special assumptions about such a process.

We model contests as reinforcement-learning processes, in which contestants update estimated values of performing an aggressive action A compared to a submissive action S. We use a simple form of reinforcement learning in which the action value is updated with a learning rate [26], and an individual chooses the aggressive action in a round of a contest if the estimated value is positive (there is also a small probability of using a random action). There are other variants of reinforcement learning that can be used to model contests, such as actor-critic leaning [4, 5]. Here we use the simpler action-value learning, in order to reduce the complexities of coalitional contests somewhat.

**Table S1:** Definitions and notation for the model.

| notation | definition or explanation |
| --- | --- |
| A, S | available actions: A is aggressive, S is submissive |
| $q_i$ | quality (fighting ability) of individual $i$ |
| $\mu_q, \sigma_q$ | mean and SD of normal distribution of quality; $\mu_q = 0$ |
| $c_{ij}$ | perceived cost by $i$ from pairwise AA round against $j$<br>given by $c_{ij} = \exp(-(q_i - a_1 q_j) + \epsilon_{ij})$; eq. (S1) |
| $a_1$ | multiplier of $q_j$ in perceived cost; $0 < a_1 \leq 1$ |
| $\epsilon_{ij}, \sigma_c$ | random influence $\epsilon_{ij}$ with SD $\sigma_c$ in perceived cost $c_{ij}$ |
| $v_{ti}, v_{bi}$ | perceived reward by $i$ of performing the aggressive action A in territorial and boundary contests (genetically determined traits) |
| $r_{ij}$ | perceived reward by $i$ from pairwise round against $j$; eq. (S2);<br>$r_{ij} = v_i - c_{ij}$ for AA round; $r_{ij} = v_i$ for AS round |
| $w_{ij}$ | estimated reward value by $i$ of using action A<br>against $j$ in the current round of a pairwise contest;<br>use action A in pairwise round against $j$ if $w_{ij} > 0$ |
| $u_{0i}, u_{di}$ | starting value of $w_{ij}$ and increment to $w_{ij}$ after coalitional dropout (genetically determined traits) |
| $\alpha_i$ | learning update rate for $i$ (genetically determined trait)<br>with pairwise update $w_{ij} \leftarrow w_{ij} + \alpha_i(r_{ij} - w_{ij})$; eqs. (S3, S4) |
| $C_{ijk}$ | perceived cost by $i$ from coalitional fighting round against $j, k$;<br>$C_{ijk} = b_1 [\exp(-(q_i - a_1 q_j) + \epsilon_{ij}) + \exp(-(q_i - a_1 q_k) + \epsilon_{ik})]$; eq. (S7) |
| $C_{ji}, C_{ki}$ | perceived cost by $j$ or $k$ from coalitional fighting round against $i$;<br>$C_{ji} = b_2 \exp(-(q_j - a_1 q_i) + \epsilon_{ji})$; eq. (S8) |
| $R_{ijk}$ | perceived reward by $i$ from coalitional round against $j, k$;<br>$R_{ijk} = v_i - C_{ijk}$ for AAA round; eq. (S9) |
| $R_{ji}, R_{ki}$ | perceived reward by $j$ or $k$ from coalitional fighting round against $i$;<br>$R_{ji} = v_j - C_{ji}$; eq. (S9) |
| $W_{ijk}, W_{ji}, W_{ki}$ | role-specific coalitional estimated action values; eq. (S10, S11, S12) |
| $U_{1i}, U_{2i}, U_{3i}$ | increments for the starting action values in a coalitional contest |
| $V_i, V_j, V_k$ | role-specific resource values for winning coalitional contest |
| $V_t, V_b$ | resource values for territorial and boundary contests |
| $L_k$ | helping liability: individual $k$ intervenes if $L_k > 0$; eq. (S6) |
| $h_{0k}, h_{1k}, h_{2k}$ | parameters for helping liability (genetically determined traits) |
| $C_h$ | opportunity cost of helping: $C_h = 0.02$ |
| $D_i$ | accumulated fighting damage for $i$; each fighting round between $i$ and $j$ increases damage by $c_{ij}, C_{ijk}$ etc.; eqs. (S5, S13) |
| $A_i$ | accumulated reproductive resources; eqs. (S14, S15) |

The overall sequence of events, corresponding to the lifetimes of the challenger *i*, the defender *j*, and the neighbour *k*, are then as follows. There is a border contest between *j* and *k*, and some time later *i* challenges *j* for ownership, and then *k* decides whether to intervene, after which there is either a coalitional or a pairwise contest. If *i* wins there is also a border contest between *i* and *k*. From these contests, *i*, *j*, and *k* accumulate fitness increments corresponding to the contested resource values if they win, and they also accumulate fighting damage. For the remainder of their lives, we assume that there is one additional unit of fitness increment available to an undamaged individual. With contest damage, this is multiplied by a factor like exp(*−d*_1_*D_i_*), where *D_i_* is the accumulated damage for *i* (and similarly for *j* and *k*), and *d*_1_ is a parameter that determines how dangerous the fighting is in terms of potential fitness losses.

#### Pairwise contests

Our modelling of pairwise contests is inspired by action-value reinforcement learning [26], which can be regarded as Sarsa learning when the states for decision-making describe whether the contest is ongoing or not. There are two contestants, here denoted *i* and *j* with fighting abilities *q_i_* and *q_j_*, and the contest consists of a sequence of rounds. The fighting abilities are drawn from a normal distribution with mean 0 and standard deviation *σ_q_*. Pairwise contests between *j* and *k*, or between *i* and *k*, are implemented similarly, except for changes in the contested resource value. In each round, a contestant can use either an aggressive action A or a submissive action S. Individual *i* maintains a variable *w_ij_*, interpreted as an estimated value of using the aggressive action A in a round of interaction with *j*, and similarly for other individuals. The value of using the submissive action S is assumed to be zero, so *w_ij_* can be interpreted as the estimated value of the difference between using A and using S. With a small probability (we used 0.001) an individual uses a random action and otherwise uses action A if the estimated value is positive. This is sometimes referred to as an *ɛ*-greedy policy [26].

At the start of the life of individual *i* we have *w_ij_* = *u*_0_*_i_*, where *u*_0_*_i_* is an evolving trait, and similarly for the starting values for other individuals. We assume that an individual *i* perceives a reward *v*_t_*_i_* from performing A in a territorial contests, and similarly *v*_b_*_i_* in a border contest, and that these are evolving traits, whereas the perceived reward from performing S is zero. The reason for introducing two traits is that a contest over territory ownership is for a higher resource value than a border contest. There are also perceived costs (penalties) from a fighting round, i.e., an AA round. With fighting abilities *q_i_* and *q_j_*, the perceived costs are

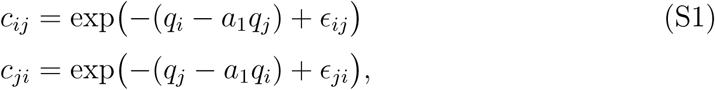

where the *ɛ_ij_* and *ɛ_ji_* are independent normally distributed errors of observation with mean zero and standard deviation *σ_c_*, and *a*_1_ is a factor giving the type of assessment. For *a*_1_ = 1 there is mutual assessment, in the sense that costs depend only on the difference in fighting abilities, and we refer to this as unbiased costs. For 0 *< a*_1_ *<* 1 there is an element of self assessment, which we refer to a self-biased costs: perceived costs depend more strongly on own fighting ability *q_i_* than opponent fighting ability *q_j_*. The net perceived rewards from an AA round in a territorial contest are then

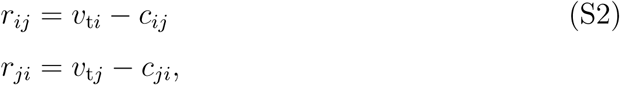

and similarly for a border contest. We have *r_ij_* = *v*_t_*_i_* and *r_ji_* = 0 for an AS round, *r_ij_* = 0 and *r_ji_* = *v*_t_*_j_* for an SA round, and *r_ij_* = *r_ji_* = 0 for an SS round.

The *w_ij_* and *w_ji_* are estimates of rewards from a round where the individual, *i* or *j*, uses the action A, and for these we have the so-called prediction errors *δ_ij_* = *r_ij_ − w_ij_* and *δ_ji_* = *r_ji_ − w_ji_*. The prediction errors are used for reinforcement learning updates, as follows. If *i* uses action A in a round, the update is

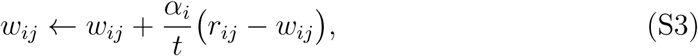

where the learning rate *α_i_* is an evolving trait of individual *i* and *t* is the round of the contest (*t* = 1*, . . .*). For a round where *i* uses S, the estimate *w_ij_* is unchanged. Similarly, for *j* we have

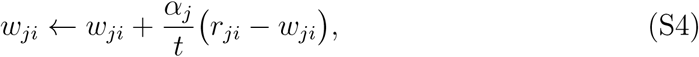

for a round where *j* uses action A. In addition, for a pairwise contest over territory ownership, such as when the defender or the neighbour drops out of a coalitional contest, or if the neighbour does not intervene, an amount *u*_d_*_i_* is added to *w_ij_* or *w_ik_*, and similarly for *w_ji_* or *w_ki_*, where *u*_d_*_i_* is an evolving trait of *i*.

These assumptions mean that a pairwise contest is represented as realizations of two stochastic processes, *w_ij_* and *w_ji_*. A contest ends either after two AS rounds in a row, with *i* as winner, or two SA rounds in a row, with *j* as winner, or after a single SS round, which is a draw. Requiring two rounds in a row reduces the chance that one individual ‘accidentally’ gives up (in practice this requirement does not make a big difference). See Fig. 1A,B,C and Fig. 2A,B,C,F in the main text for illustrations of perceived costs, trajectories of action value estimates, and probabilities of winning and durations of contests as functions of the fighting abilities *q_i_* and *q_i_*.

There is an accumulation of damage from the AA rounds in a pairwise contest.

For individual *i* we have

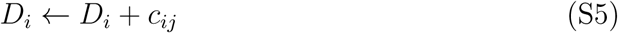

from a pairwise fighting round, with perceived cost *c_ij_* from eq. (S1). There is a similar expression for the damage *D_j_*. The total damage an individual accumulates leads to a fitness cost, described by eq. (S15) below.

#### Decisions to intervene

The neighbour *k* bases the decision of whether to intervene on a threshold condition that a helping liability *L_k_* is positive. The helping liability is a linear function of the action-value estimates *w_ki_* and *w_kj_*, and the condition is that

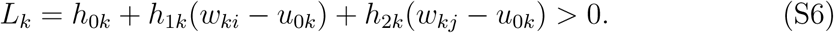

The parameters *h*_0_*_k_*, *h*_1_*_k_*, and *h*_2_*_k_* are genetically determined traits, and the variables *w_ki_ − u*_0_*_k_* and *w_kj_ − u*_0_*_k_* are deviations of the action-value estimates from their initial values. These variables will be positively correlated with the fighting ability differences *q_k_ − q_i_* and *q_k_ − q_j_*. For instance, our evolutionary simulations show that the coefficient *h*_2_*_k_* evolves to become positive and larger than *h*_1_*_k_*, suggesting that a main explanation for intervention is that it is an attempt by the neighbour *k* to help a weaker defender *j*to defeat a challenger *i*, who otherwise could take over the territory. See Figures 3A,D, 4A,D, and 5A,D in the main text for illustrations of this.

#### Coalitional contests

Our model of coalitional contests is similar to that for pairwise contests, in that individuals update an estimate of the reward from using the action A, but the expressions for the perceived costs differ. For the challenger, the perceived cost *C_ijk_* of using A in round where the coalition members *j* and *k* also use A depends on the fighting abilities *q_i_*, *q_j_*, and *q_k_* and is given by

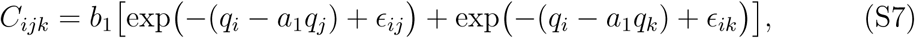

where the *ɛ_ij_* and *ɛ_ik_* have normal distributions with an SD of *σ*_c_, *a*_1_ is the same factor as in eq. (S1), and *b*_1_ is a parameter that sets the strength of the effect of the combined aggression from the coalition members (we sometimes use *b*_1_ = 1). For the coalition members, the perceived costs are

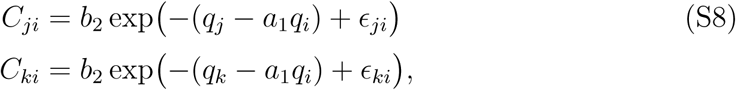

where *b*_2_ is a parameter that sets how much the aggression by *i* is weakened when fighting two simultaneous opponents (we sometimes use *b*_2_ = 0.5). The perceived rewards of a coalitional fighting round are then

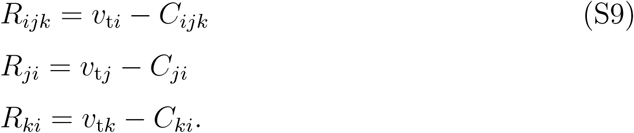

We use the notation *W_ijk_*, *W_ji_*, and *W_ki_* for the estimates by the contestants of these rewards. At the start of a coalitional contest we assume that *i*, *j*, and *k* take into account both any previous contest and their role in the coming contest to give starting values for these estimates. We assume that

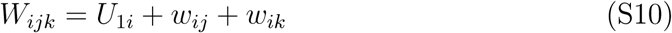

is the starting estimate for *i*, where *U*_1_*_i_* is an evolving trait of the challenger *i*. For the coalition members we have the starting values

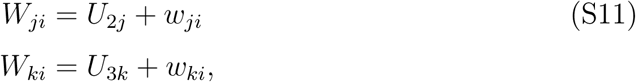

where *U*_2_*_j_* and *U*_3_*_k_* are evolving traits of respectively the defender *j* and the neighbour *k*. The values of the traits *U*_1_*_i_*, *U*_2_*_j_* and *U*_2_*_k_* influence how persistent in fighting an individual will be in a given role in a coalitional contest. For instance, a negative *U*_1_*_i_* causes a challenger to give up more readily when meeting two opponents in a coalitional contest, and positive *U*_2_*_j_* and *U*_3_*_k_* cause the coalition members to not give up readily. Because rewards of choosing the action A are at most the perceived value, for instance *v*_t_*_i_* for *i*, we impose the constraint that the estimates *W_ijk_*, *W_ji_*, and *W_ki_* cannot exceed this value. The learning updates from a coalitional fighting round are then

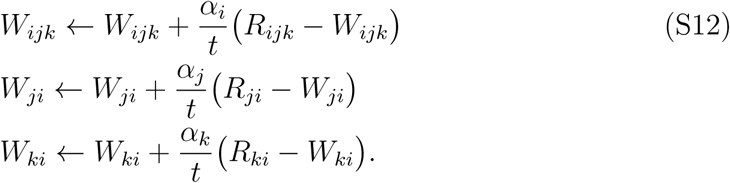

Just as for pairwise contests, there is an accumulation of damage from coalitional fighting rounds. We have

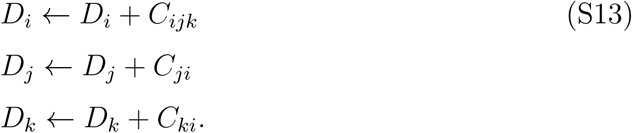

Again, the total damage an individual accumulates leads to a fitness cost, described by eq. (S15) below.

#### Fitness costs and benefits from a contest

Concerning reproduction we assume than an individual accumulates reproductive resources, including mating success, during its life, and that the individual’s reproductive success is proportional to the accumulated resources. So, if *A_i_* is the accumulated resources for *i*, and *i* wins a contest with resource value *V_i_*, we have

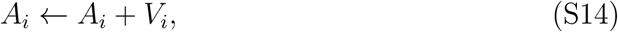

with similar expressions for cases with other resource values. In addition to resources from winning the contests described here, we assume that there is a remaining period of the life of an individual where reproductive resources are accumulated, and that contests damage influences how much of these resources an individual can acquire, resulting in

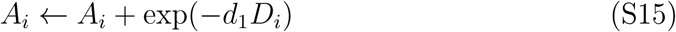

for individual *i*, where *D_i_* is the accumulated fighting damage for *i*, as described in eqs. (S5, S13), and *d*_1_ is a parameter setting the strength of the fitness cost of damage. There are similar expressions for *j* and *k*. Note that we assume proportionality between accumulated perceived costs (the perceived damage *D_i_*) and the effect of contest damage on fitness (*d*_1_*D_i_* in eq. S15). This need not be the case and different assumptions could influence the evolution of fighting behaviour, but we have not further investigated this issue.

#### Evolving traits and evolutionary simulations

Each individual has a number of genetically determined traits that can evolve in the simulations. So, for individual *i*, there are traits for the starting action-value estimate *u*_0_*_i_*, the increment *u*_1_*_i_* for pairwise ownership contests, the increments *U*_1_*_i_*, *U*_2_*_i_*, and *U*_3_*_i_* for the starting action-value estimates in coalitional contests in eqs. (S10, S11), the learning rate *α_i_* in eqs. (S3, S4, S12), and the parameters *h*_0_*_i_*, *h*_1_*_i_*, and *h*_2_*_i_* in the expression for the helping liability *L_i_* in eq. (S6), controlling the intervention decision when in the role of neighbour. In our simulations, individuals have an additional trait, in the form of a generalization factor determining how information from a coalitional contest influences subsequent pairwise contests, but these traits do not play any important role for our results, so we do not describe them in any detail here.

A simulated population consists of 2000 groups of three individuals, assumed to be hermaphrodites, with the genetics of each trait being a breeding value of an independent implementation of the infinitesimal model of quantitative genetics [28], making up a total population of 6000 individuals. The standard deviation of segregation deviations at reproduction for each trait is adjusted to correspond to the range of trait variation, to ensure that simulations could locate evolutionary equilibria.

When producing an offspring for the next generation, parents are selected with probability proportional to the accumulated resources described in eqs. (S14, S15), over the entire set of groups. In this way there are no effects of relatedness between interacting individuals.

Simulations were first run for at least 10^6^ generations to reach a starting point of approximate equilibrium, and then continued for another 2 *×* 10^6^ generations. The mean trait values over these 2 *×* 10^6^ generations were used to generate the results shown in figures, together with the means of the genetic variation for each trait. We also used the mean of the SD of each trait over 2 *×* 10^6^ generations to generate genetic variation for the results. For cases where evolution led to neighbours never intervening, we reduced this to 10^6^ generations.

A reason for the many generations of simulation is that, for a multi-trait evolutionary equilibrium, stabilizing selection can be relatively weak for some of the traits. The equilibrium outcome is then that traits fluctuate over time, with notable temporal autocorrelation. We illustrate this phenomenon in Fig. S1.

**Figure S1:**
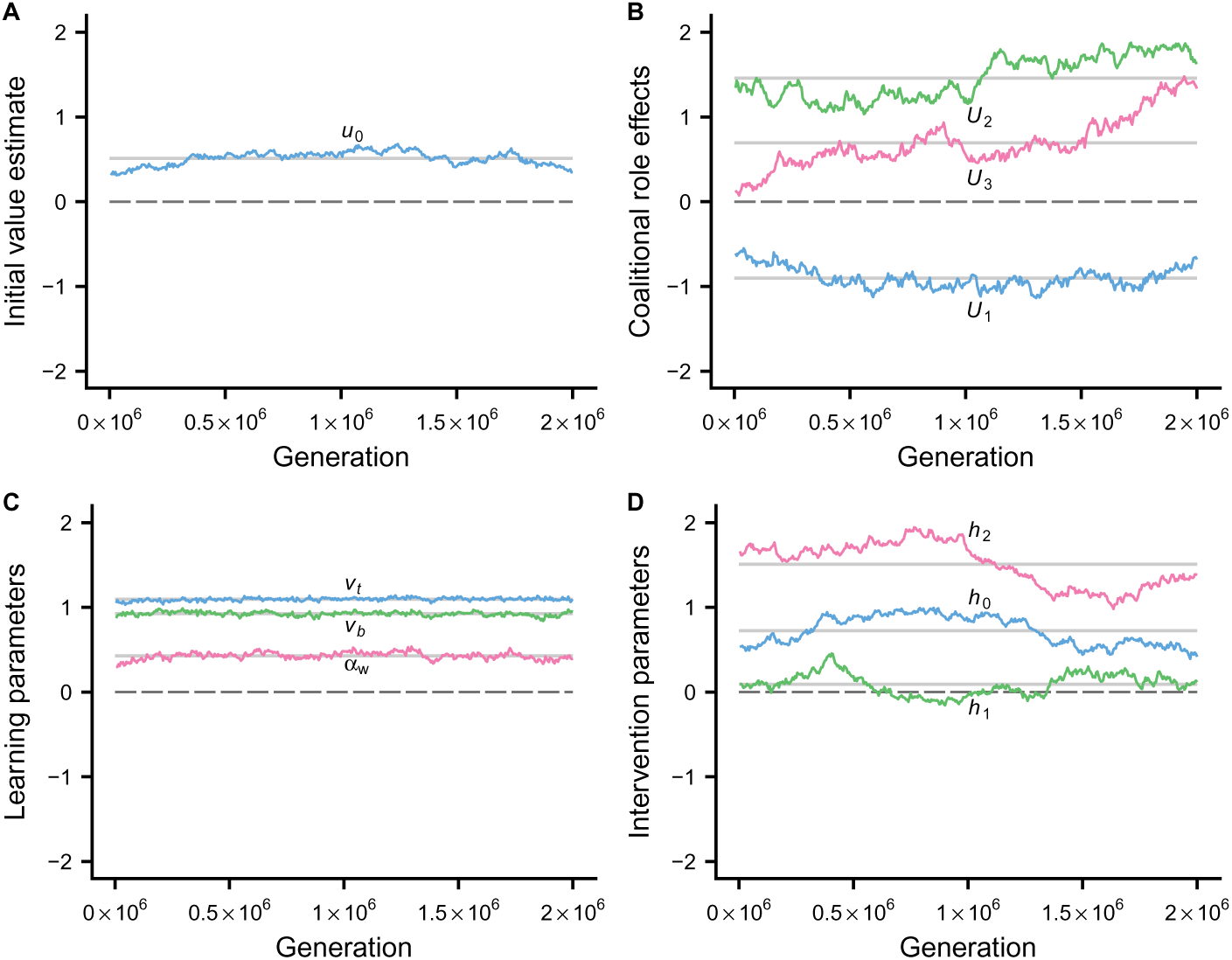
Illustration of evolutionary trajectories corresponding to the simulations in Fig. 2.

### Comparison of action-value learning and sequential assessment

Results from the reinforcement-learning representation of pairwise contests shows some similarity to those from the sequential assessment model Enquist and Leimar [27], as illustrated for action-value learning in Figs. S2, S3. As discussed by McNamara and Leimar [4], the original sequential assessment game is a so-called small-worlds model with an evolutionarily stable switching line that is an ESS that properly accounts for Bayesian updating of the information available to an individual (given the assumption that *q_i_* and *q_j_* are randomly drawn from some prior distribution). In comparison, models with reinforcement learning are instead large-worlds models [4], with the evolutionary outcome an approximate multi-trait equilibrium.

**Figure S2:**
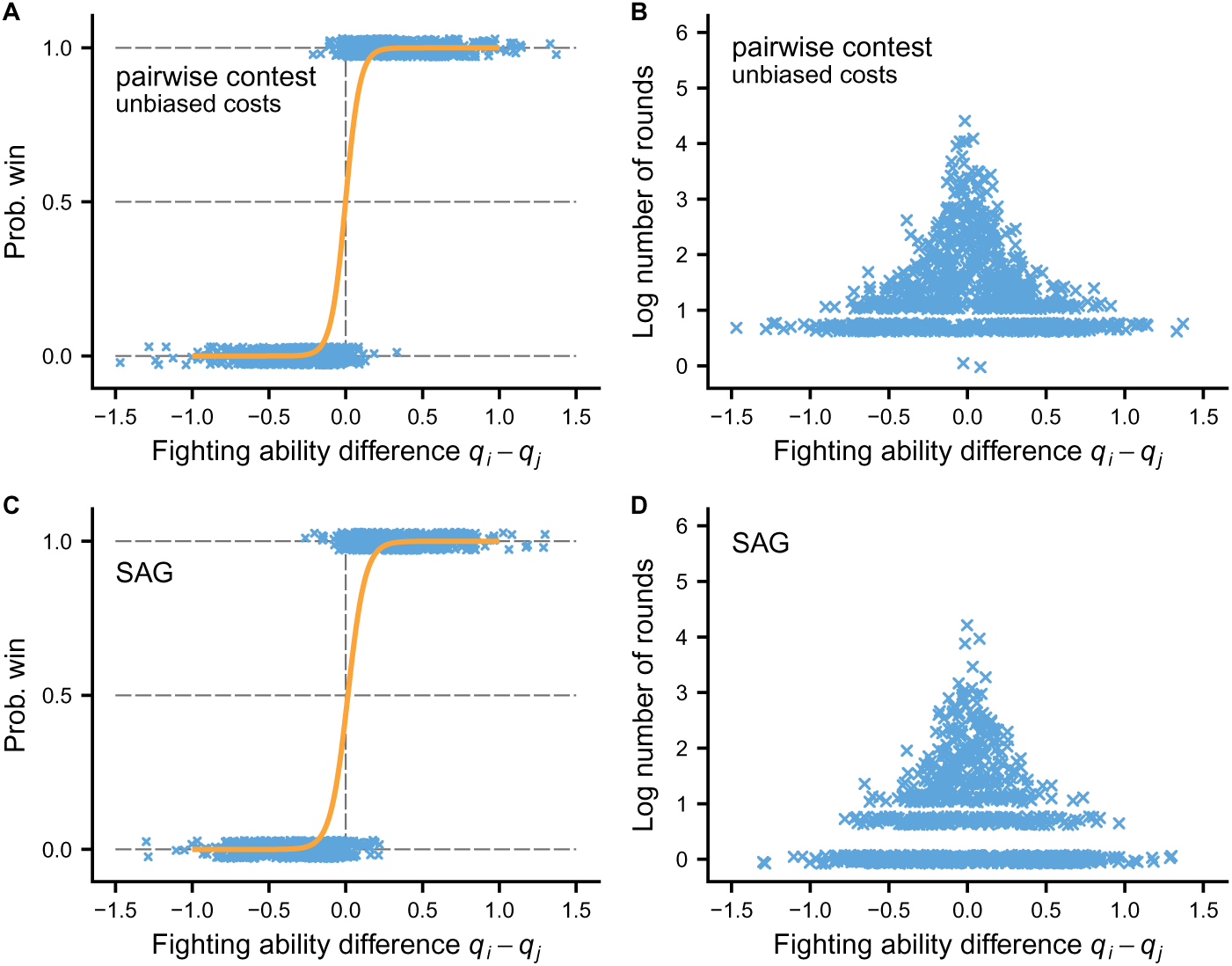
Comparison of pairwise action-value learning contests with the sequential assessment game (SAG), for border conflicts. The points in the panels illustrate probabilities of winning and durations for 1000 contests, as a function of the difference *q_i_ − q_j_* in fighting ability between opponents. For action-value learning, the perceived costs are as shown in Fig. 1A in the main text (unbiased costs), and the contested resource value, *V* = *V*_b_ = 0.25, corresponds to boundary contests in the presentation in the main text, which are illustrated in Fig. 2F. The parameters for the sequential assessment game are chosen to approximately correspond to this situation. Comparing panels (A) and (C) shows high similarity in the probabilities of winning, whereas comparing (B) and (D) shows some differences in the distributions of contests durations. Compared to the sequential assessment game, action-value learning has fewer contests that end directly, without any fighting (bottom-most points in the panels), but the overall durations are still fairly similar.

**Figure S3:**
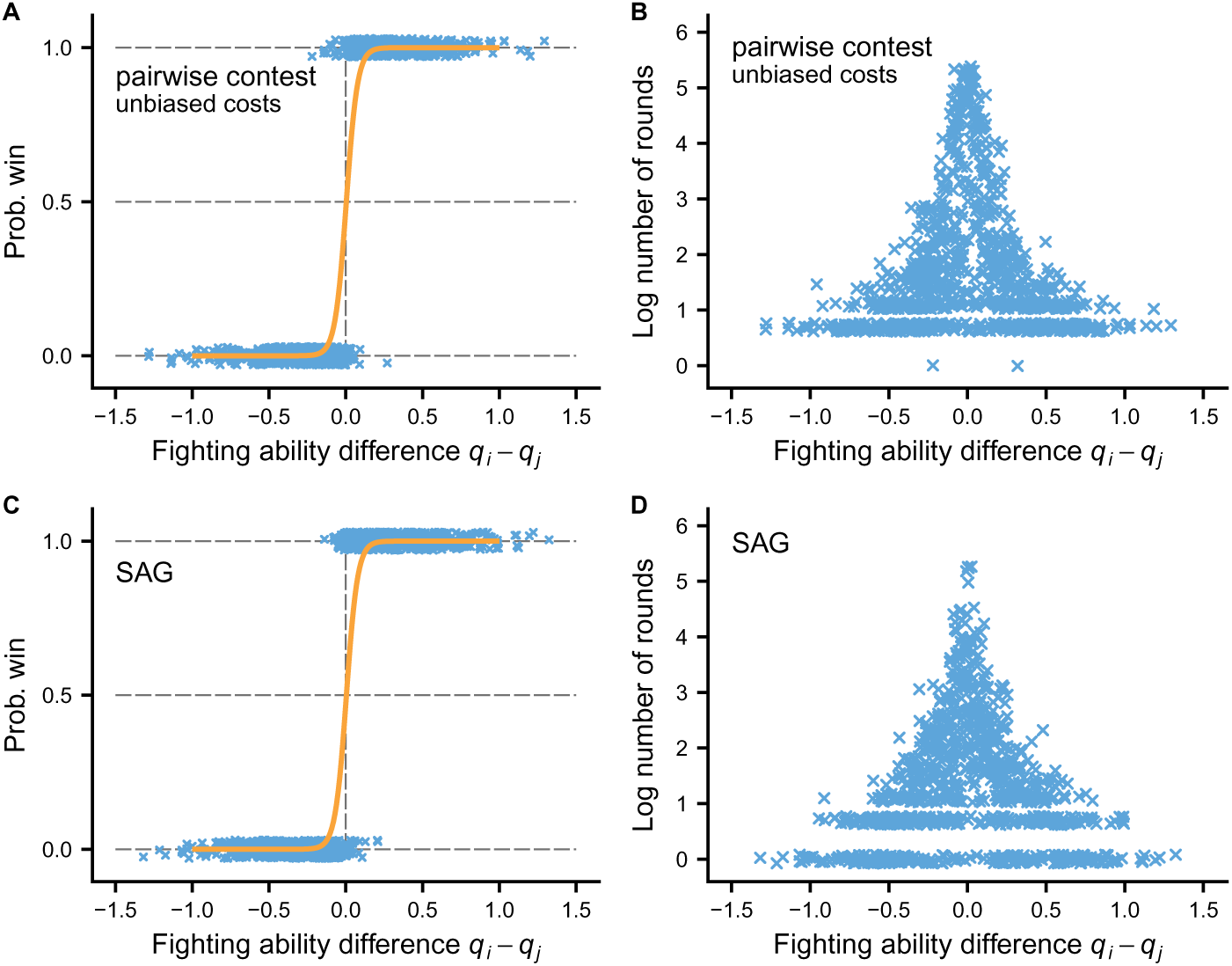
Comparison of pairwise action-value learning contests with the sequential assessment game (SAG), for territorial conflicts. The points in the panels illustrate probabilities of winning and durations for 1000 contests, as a function of the difference *q_i_ − q_j_* in fighting ability between opponents. For action-value learning, the perceived costs are as shown in Fig. 1A in the main text (unbiased costs), and the contested resource value, *V* = *V*_t_ = 1.0, corresponds to pairwise territorial contests in the presentation in the main text, which are illustrated in Fig. 4E,F and Fig. 5E,F. The parameters for the sequential assessment game are chosen to approximately correspond to this situation. Because of a higher resource value, the contests tend to be longer here than in Fig. S2 and, as a consequence, winning is determined more sharply by relative fighting ability, both for action value learning in (A) and sequential assessment in (C). Comparing (B) and (D) again shows some differences in the distributions of contests durations, with action-value learning having fewer contests that end directly, without any fighting (bottommost points in the panels), but the overall durations are fairly similar.

### Comparison with actor-critic learning

Here we illustrate that our implementation of action-value learning leads to similar results as actor-critic learning for pairwise contests (see Fig. S4). Previous work on modelling dominance hierarchies have used actor-critic learning as a behavioural mechanism [4, 5, 6, 7]. A characteristic of this form of contest model is that individuals can show rapid shifts from being aggressive to being submissive (see, e.g., Fig. 2 in [5]), which is a consequence of the actor learning rate evolving to a high value. This is qualitatively similar to action-value learning when the estimated value shifts from positive to negative.

**Figure S4:**
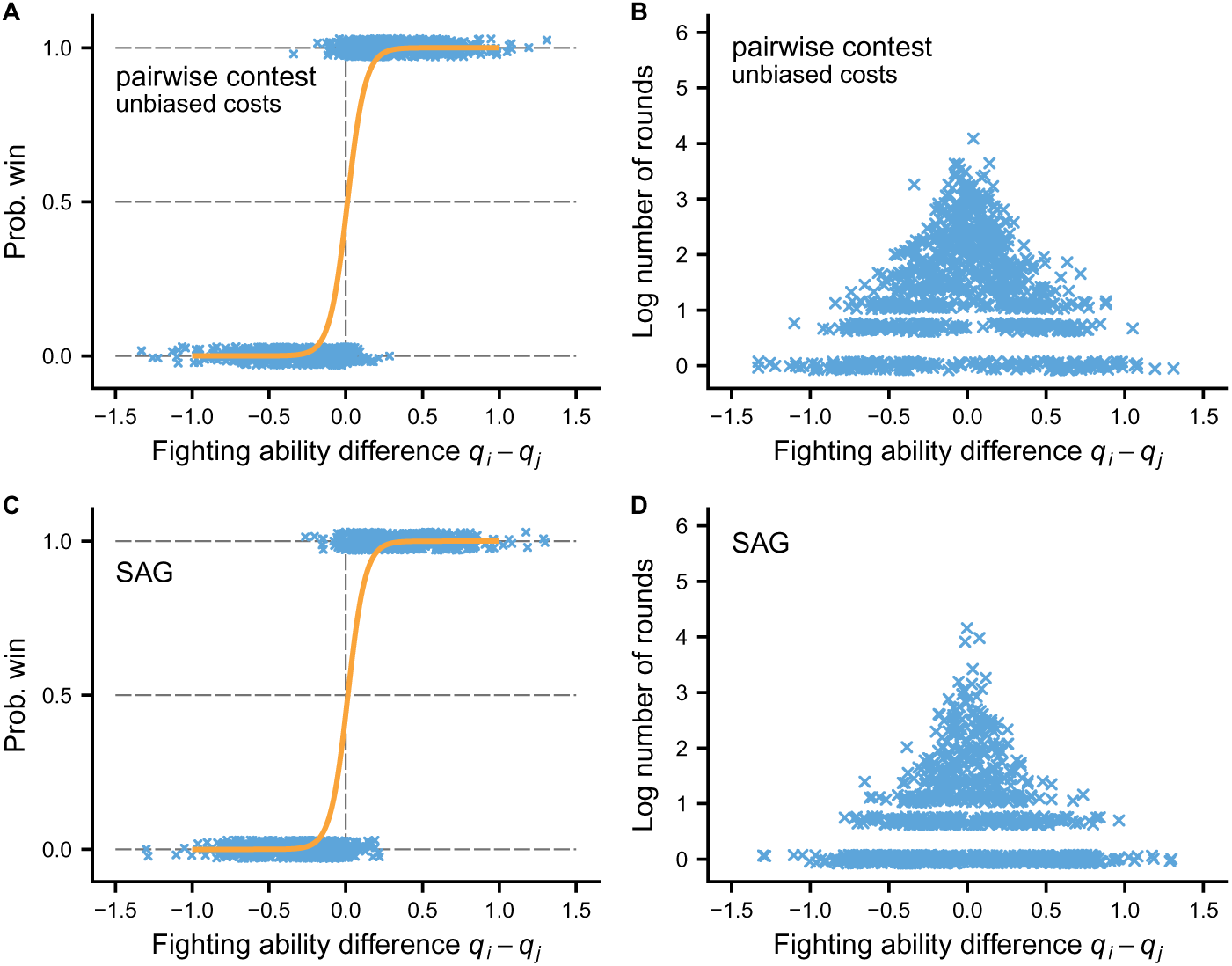
Comparison of pairwise actor-critic learning contests with the sequential assessment game (SAG), for a situation corresponding to boundary conflicts with unbiased cost in the presentation in the main text. The contested resource value is *V* = *V*_b_ = 0.25. The points in the panels illustrate probabilities of winning and durations for 1000 contests, as a function of the difference *q_i_ − q_j_* in fighting ability between opponents. Comparing panels (A) and (C) shows high similarity in the probabilities of winning, just as for action-value learning. Further, comparing (B) and (D) shows a somewhat better correspondence in the distribution of durations than is the case for action-value learning in Fig. S2.

Actor-critic learning portrays a decision maker as being divided into two parts, an actor and a critic (see chapter 15 of [26] for a presentation of this perspective), which means that more evolving traits are needed for an actor-critic mechanism. In particular, for such a mechanism to learn about fighting abilities in a contest, some form of observation of fighting abilities need to be represented in the action preferences of the actor; it is not enough that the fighting abilities influence the perceived rewards, as is the case for the action-value learning model we present here. Examples of how such observations can be implemented in actor-critic learning are found in chapter 8 of [4] and in [5].

### Effects of biases in perceived costs

Differences in how perceived costs depend on fighting abilities, for instance from having different values of the parameter *a*_1_ (eqs. S1, S7, S8; see also Figs. 1, 2 in the main text), can influence how the probability of winning and the contest duration at an evolutionary equilibrium depend on the fighting abilities. Inspired by Arnott and Elwood [8], including their Figure 1, it has become common to examine such differences in a type of diagnostic plots, in which contest duration is displayed separately as a function of the winner fighting ability and the loser fighting ability. This is illustrated for strongly self-biased costs in Fig. S5C,D, showing increasing duration both as a function of winner and loser fighting ability, but with a steeper increase as a function of loser fighting ability (Fig. S5D; the model used here has an additional trait, reducing the value *v_i_* of performing the action A as the contest gets longer, in this way reducing the length of contests). This is then diagnostic of strongly self-biased costs. The same kind of diagnostic plots for moderately self-biased and unbiased costs are shown in Fig. S6.

**Figure S5:**
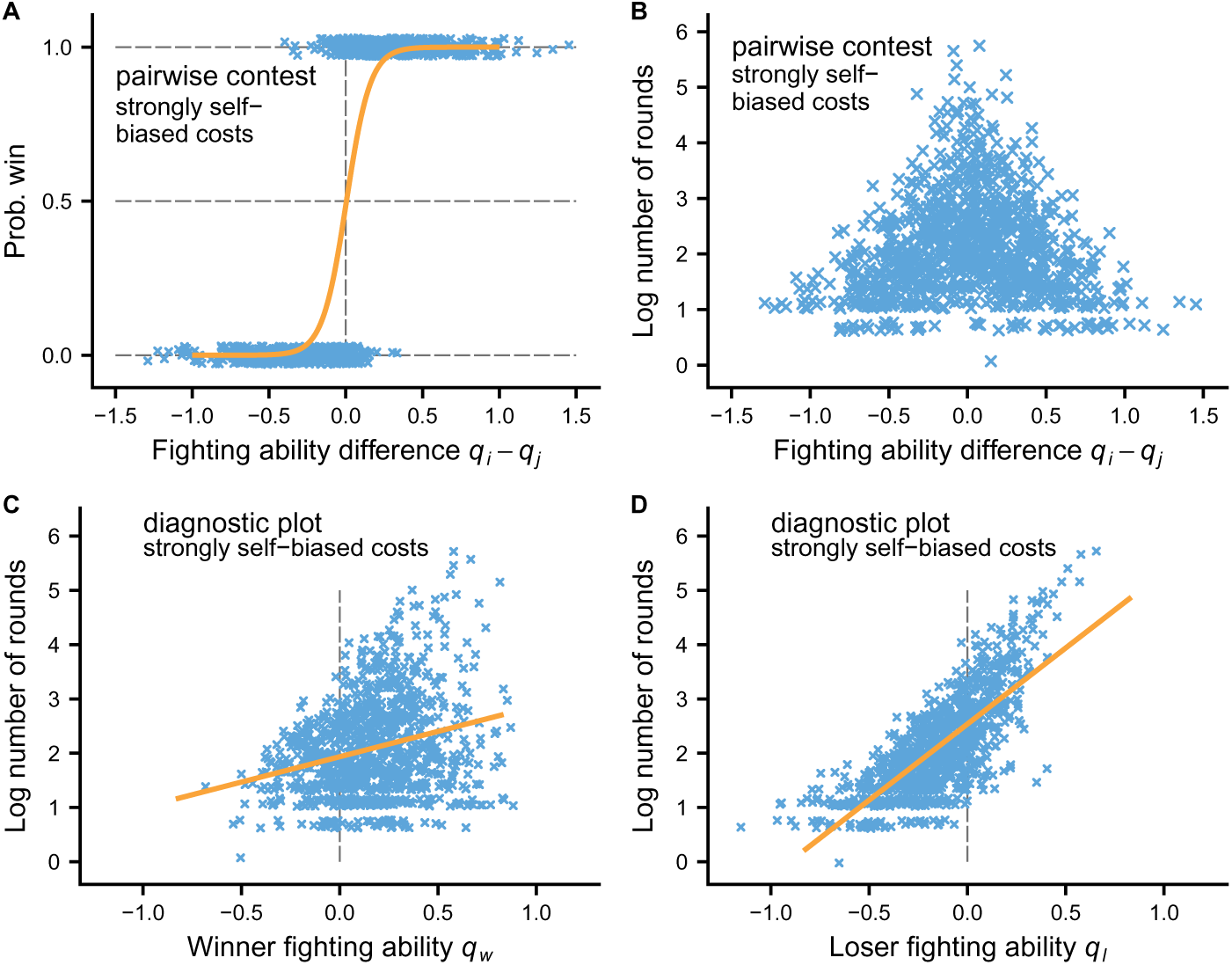
Illustration of the evolutionary outcome for an action-value learning model of pairwise contests with strongly self-biased costs (parameter *a*_1_ = 0.25 in eq. S1), together with diagnostic plots as suggested by Arnott and Elwood [8]. Note that the probability of winning is not so sharply determined by the difference in fighting ability (A) and that contests where both winner and loser have high fighting ability can become very long.

**Figure S6:**
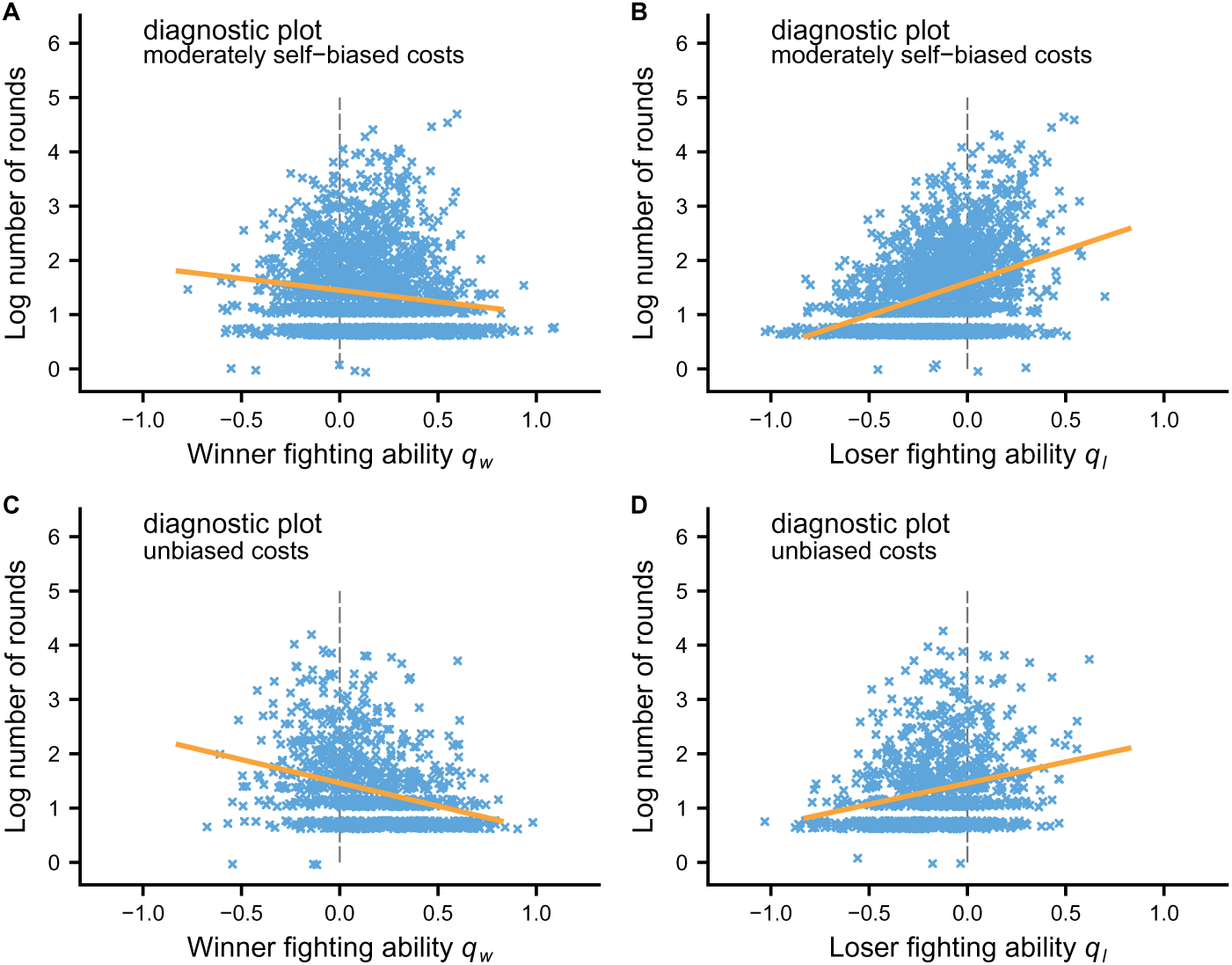
Similar diagnostic plots as in Fig. S5D,E, but here for moderately self-biased costs (*a*_1_ = 0.85) vs. unbiased costs (*a*_1_ = 1.0). For both these cases, contest duration tends to decrease with winner fighting ability and to increase with looser fighting ability.

**Figure S7:**
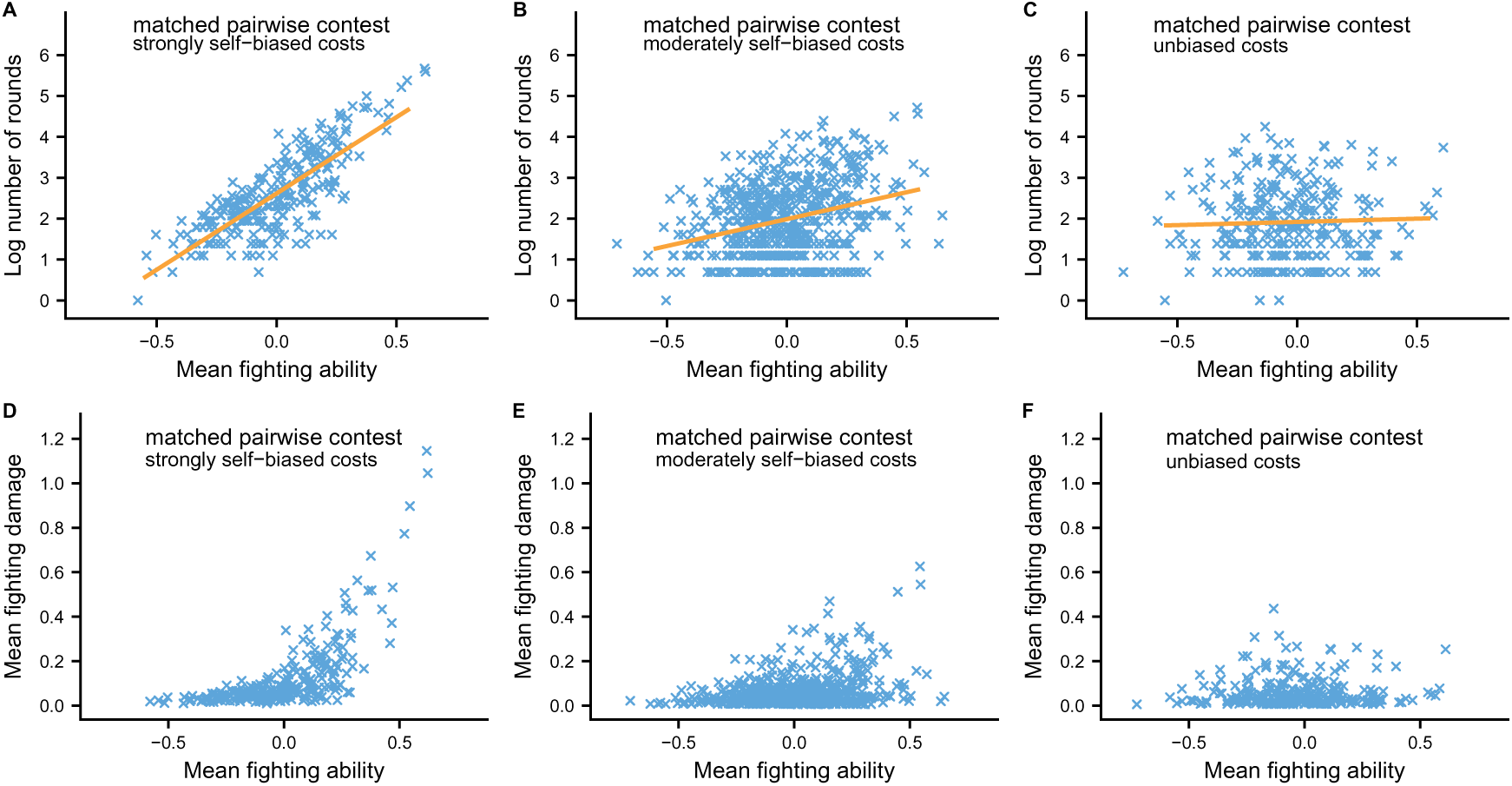
Alternative diagnostic plots based on the duration of contests between approximately matched individuals (*|q_i_ − q_j_| <* 0.15) as a function of the mean fighting ability, together with the accumulated fighting damage from these contests (*a*_1_ = 0.25, 0.85, 1.0 in A and D, B and E, and C and F). Note the long contests in panels A,D.

For model simulations, and also for experiments, an alternative diagnostic is to examine contest duration as a function of the mean fighting ability of approximately matched opponents, as is done in Fig. 2C,F in the main text. Figure S7A,B,C shows additional such plots, together with plots of the accumulated damage from contests. A striking aspect of these plots is that, for strongly self-biased costs, contests between individuals of high fighting ability can become very long and result in much damage (Fig. S7A,D). This shows that there is a certain inefficiency in assessment through strongly self-biased perceived costs. If contests between individuals of high fighting ability are common, there will be selection for assessment mechanisms that are more sensitive to relative fighting ability. This possibility is an interesting but so far little studied question.

### Different contributions of coalition members to a contest

Our model assumes that the defender and the neighbour contribute equally to a coalitional contest. This is expressed in the symmetric manner in which the coalition members appear in the perceived costs, in eqs. (S7, S8). It is of course easy to modify this assumption, for instance by introducing separate parameters *b*_1_*_j_, b*_1_*_k_* and *b*_2_*_j_, b*_2_*_j_*, with equal contribution corresponding to *b*_1_*_j_* = *b*_1_*_k_* = *b*_1_ and *b*_2_*_j_* = *b*_2_*_k_* = *b*_2_.

**Figure S8:**
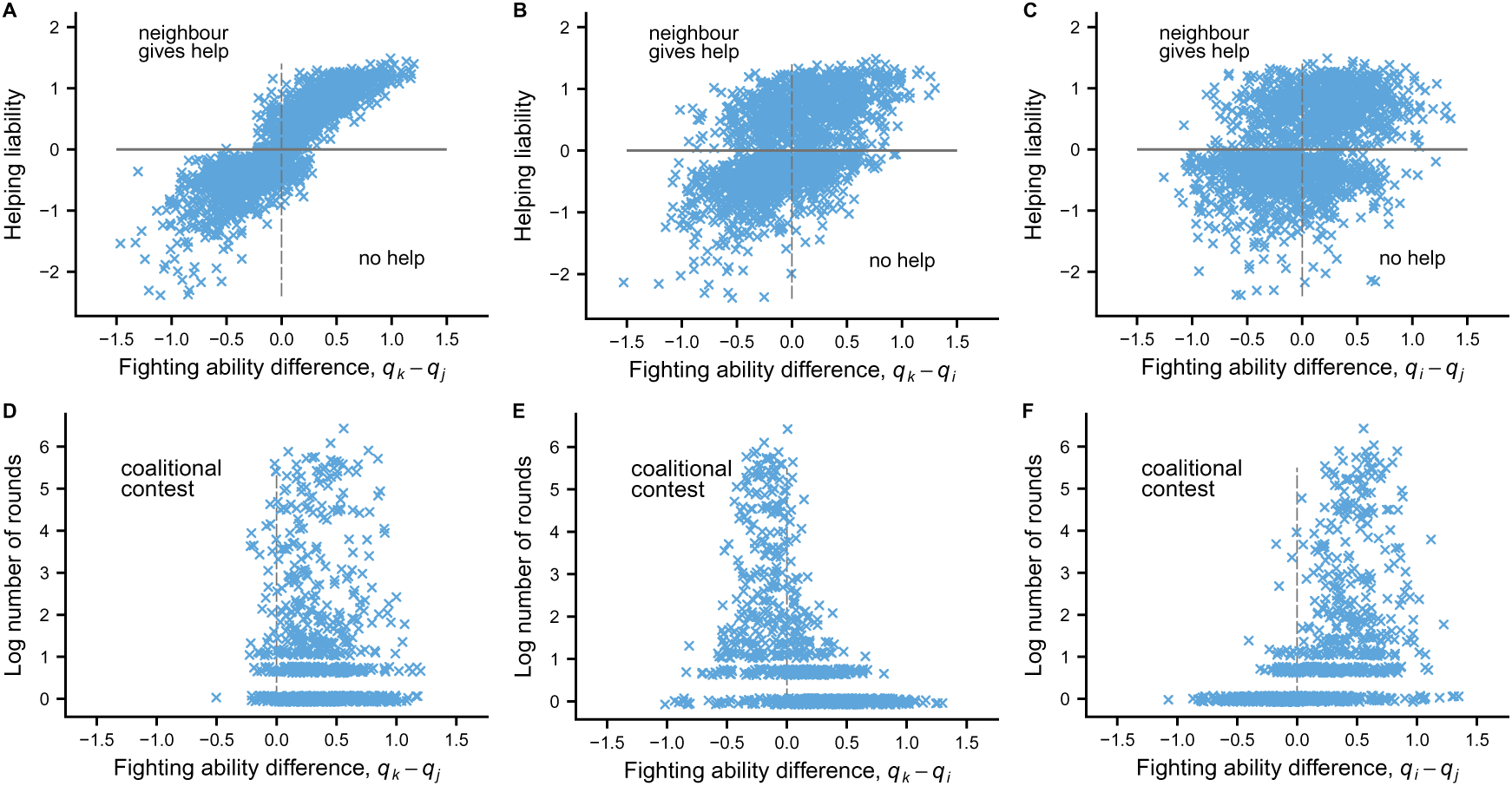
Intervention decisions by the neighbour and durations of coalitional contests in a situation where the defender contributes less than the neighbour to a coalitional contest against a challenger.

As an illustration, in Figure S8 we show results for a case where the neighbour*k* contributes more than the defender *j* to a coalition (*b*_1_*_j_* = 0.50*, b*_1_*_k_* = 0.75*, b*_2_*_j_* = 0.50*, b*_2_*_k_* = 0.75), where we also assumed that the neighbour has a bit more information about the challenger when making the decision about intervention, corresponding to one round of aggressive interaction. Overall, the results are fairly similar to those for equal contributions, as seen by comparing with Fig. 3 in the main text, but there are some differences. In Fig. S8E the longest contests, for which the coalition and the challenger are about equally matched, occur when the challenger is only somewhat stronger than the neighbour, but in Fig. 3C the challenger needs to be substantially stronger.

Concerning the effect of the challenger fighting ability on the decision by the neighbour to intervene, we see from Fig. S8B that the neighbour is somewhat more likely to intervene when having a higher fighting ability than the challenger. This would be in accordance with the observations by Detto et al. [12] on fiddler crabs.

Making the difference in contributions from coalition members more extreme, for instance by assuming the the neighbour fully takes over the contest with the challenger, results in the decision by the neighbour evolving to not intervene at all. This makes sense as it should be preferable for the neighbour to encounter the challenger in a pairwise border dispute, which would be over lower stakes than a pairwise contest over the defenders territory.

